# Phage therapy potentiates second-line antibiotic treatment against pneumonic plague

**DOI:** 10.1101/2022.02.07.479346

**Authors:** Yaron Vagima, David Gur, Moshe Aftalion, Sarit Moses, Yinon Levy, Arik Makovitzki, Tzvi Holtzman, Ziv Oren, Yaniv Segula, Ella Fatelevich, Avital Tidhar, Ayelet Zauberman, Shahar Rotem, Emanuelle Mamroud, Ida Steinberger-Levy

**Affiliations:** Department of Biochemistry and Molecular Genetics, Israel Institute for Biological Research, Ness-Ziona, Israel; Department of Biotechnology, Israel Institute for Biological Research, Ness Ziona, Israel

**Keywords:** phage therapy, antibiotic therapy, ceftriaxone, *Yersinia pestis*, plague, antibiotic resistance, ϕA1122, PST, phage-antibiotic combination

## Abstract

Plague pandemics and outbreaks have killed millions of people during the history of humankind. The disease, caused by *Yersinia pestis* bacteria, can currently be treated efficiently with antibiotics. However, in the case of multidrug-resistant (MDR) bacteria, alternative treatments are required. Bacteriophage (phage) therapy has shown efficient antibacterial activity in various experimental animal models and in human patients infected with different MDR pathogens. Herein, we evaluated the efficiency of ϕA1122 and PST phage therapy, alone or in combination with second-line antibiotics, using a well-established mouse model of pneumonic plague. Phage treatment significantly delayed mortality and limited bacterial proliferation in the lungs. However, the treatment did not prevent bacteremia, suggesting that phage efficiency may decrease in circulation. Indeed, *in vitro* phage proliferation assays indicated that blood has inhibitory effects on lytic activity, which may be the major cause of treatment inefficiency.

Combining phage therapy and second-line ceftriaxone treatment, which are individually insufficient, provided protection that led to survival of all infected animals, presenting a synergistic protective effect that represents a proof of concept for efficient combinatorial therapy in an emergency event of a plague outbreak involving MDR *Y. pestis* strains.

**Author summary:** Plague, caused by *Yersinia pestis* bacteria, can be efficiently treated with antibiotics. However, alternative therapies for the case of natively evolved or maliciously generated antibiotic-resistant *Y. pestis* must be developed. Due to the global increase in antibiotic resistance, there is renewed interest in examining the effectiveness of bacteriophage-based alternative therapies. Here, using a mouse model of pneumonic plague, we demonstrate that phage treatment significantly delayed mortality. By monitoring bioluminescence of engineered *Y. pestis* strain and live bacterial counts, we show that phage therapy effectively inhibited bacterial proliferation in the lung but not in blood. *In vitro* analyses showed decreased phage activity in the presence of blood, which probably explains the low efficacy of phage treatment alone. Because combination therapies will be used in an emergency situation, we tested the efficacy of *Y. pestis*-lysing phages as adjunctive therapy with a second-line antibiotic, ceftriaxone.

Whereas each individual treatment was insufficient, the combination provided effective protection and rescued all infected animals. These results clearly demonstrated the synergistic effect of combined phage and antibiotic therapy and represent a proof of concept for this alternative therapy against multidrug-resistant *Y. pestis* strains.

## Introduction

*Yersinia pestis*, a gram-negative bacterium and the etiological agent of plague, is a lethal pathogen that has led to three world pandemics throughout human history [1,2]. Although natural plague outbreaks are rare, plague is considered a reemerging disease [3,4]. A recent large outbreak occurred in Madagascar in 2017, where many patients presented with the pneumonic form of the disease [5,6]. Due to its lethality and the potential for person-to-person infectivity, *Y. pestis* is classified by the Centers for Disease Control and Prevention (CDC) as a tier-1 select agent [1,7]. Natural exposure of humans to the bacteria may occur through a carrier flea bite, leading to bubonic or septic plague that can be further developed into a secondary pneumonic plague, or by air transmission, leading to primary pneumonic plague [1]. Pneumonic plague is a rapidly deteriorating disease that, if not immediately treated with the recommended antibiotic, leads to death [1,3].

*Y. pestis* strains are usually susceptible to the recommended antibiotics. However, several strains isolated from rodents or from plague patients have demonstrated antibiotic resistance against first-line antibiotics such as streptomycin [3,8–11]. Moreover, a major concern exists regarding the possibility of the generation and usage of antibiotic-resistant *Y. pestis* strains in a bioterror event [3,4,7]. Currently, no licensed plague vaccine is available in Western countries [4,12], thus emphasizing the importance of developing alternative treatments such as bacteriophage (phage) therapy.

Lytic phages are bacterial viruses that demonstrate high specificity toward their hosts. Phage replication inside the host bacteria leads to bacterial lysis and cell death, releasing progeny virions in the so-called “lytic cycle”. The potential of using phages for antimicrobial therapy has been known for more than a century, since phages were independently identified by Frederik Twort (in 1915) and Felix d’Herelle (in 1917) [13]. Their advantages of selective and efficient bacterial killing, self-replication in the presence of the host, safety as treatments and simple and inexpensive preparation led to their application in the 1920s as a treatment against various infectious diseases, such as dysentery and cholera [13,14]. However, the appearance of commercialized antibiotics in the late 1940s changed the paradigm, and Western countries preferred the usage of antibiotics, which are characterized chemical molecules, over the usage of phage, a biological viral entity [13]. In contrast to the situation in the West, phage therapy continued to serve as an acceptable remedy in the former USSR, Georgia and Eastern European countries [15]. Recently, as antibiotic resistance has become a global threat to public health, renewed interest in phage therapy has emerged in Western countries. Phage efficiency and safety have been investigated using various bacterial-infected animal models and by conducting human clinical trials [16]. Moreover, an increasing number of case reports have described successful compassionate-use treatment of multidrug-resistant (MDR) bacterial-infected patients by administration of antibiotics in combination with lytic phages, selected in a personalized manner to lyse the isolated MDR pathogen (see [17,18] and recent reviews [16,19]).

In the present study, we evaluated the potential of phage therapy against pneumonic plague using a well-established mouse model. The therapeutic potential of treatment with the *Y. pestis*-specific lytic phages ϕA1122 and PST was assessed alone, as a dual cocktail, or in combination with the second-line broad-spectrum cephalosporin antibiotic ceftriaxone.

## Results

### Intranasal administration of ϕA1122 leads to high phage titers in the lungs

Pneumonic plague involves *Y. pestis* dissemination from the lung to internal organs and blood [20]. Thus, for efficient phage treatment, phage distribution in those tissues is a prerequisite. Therefore, we tested the tissue distribution of the *Y. pestis*-specific lytic phage ϕA1122 in C57BL/6J mice and examined its dependency on the administration route. Mice were inoculated intranasally (IN) or intraperitoneally (IP) with a single dose of phage suspension (1×10^9^ plaque-forming units (PFU)/mouse). As shown in Fig 1, phages reached all tested tissues (blood, lung, spleen and liver) within half an hour. The IN route seemed advantageous for the treatment of pneumonic plague, as it led to the highest phage concentration in the lung (∼1×10^9^ PFU/lungs, Fig 1B). Moreover, high phage concentrations were maintained in the lungs for at least 4 days (Fig 1B). Thus, we further tested the potential of intranasal treatment with ϕA1122 against *Y. pestis* airway infection.

**Fig 1.**
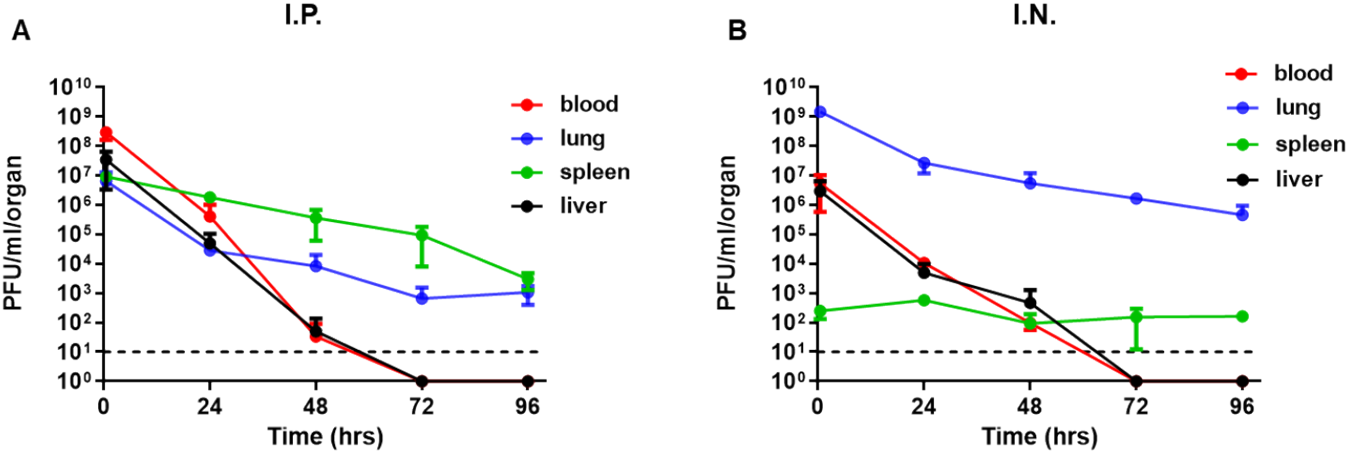
Pharmacokinetic analysis of ϕA1122 in naïve mice. A single dose of ϕA1122 phage suspension (1×10^9^ PFU) was administered to naive C57BL/6J mice, either by (A) IP injection (0.5 ml) or (B) via the IN route (35 μl). For each administration route, n=3 for each time point. Phage titration was performed using a spot assay test. Each dot represents the mean value in terms of PFU/organ (lung – blue dots, spleen – green dots, liver – black dots) or PFU/ml blood (red dots). Bars represent the standard deviation (SD).

### Intranasal administration of a single dose of phage suspension delayed mortality in a mouse model of pneumonic plague

To evaluate the benefit of phage therapy, we IN administered a single dose of the ϕA1122 lytic phage (1×10^9^ PFU/mouse) at 5 h post IN infection with a lethal dose of 10xLD_50_ *Y. pestis* strain Kim53.

As shown in Fig 2A, *Y. pestis*-infected control mice died within 3-4 days post infection (pi), showing a mean time to death (MTTD) of 3 days. In contrast, phage-treated mice showed a significant delay in time to death, which occurred within 4-7 days pi, with a MTTD of 5 days.

**Fig 2.**
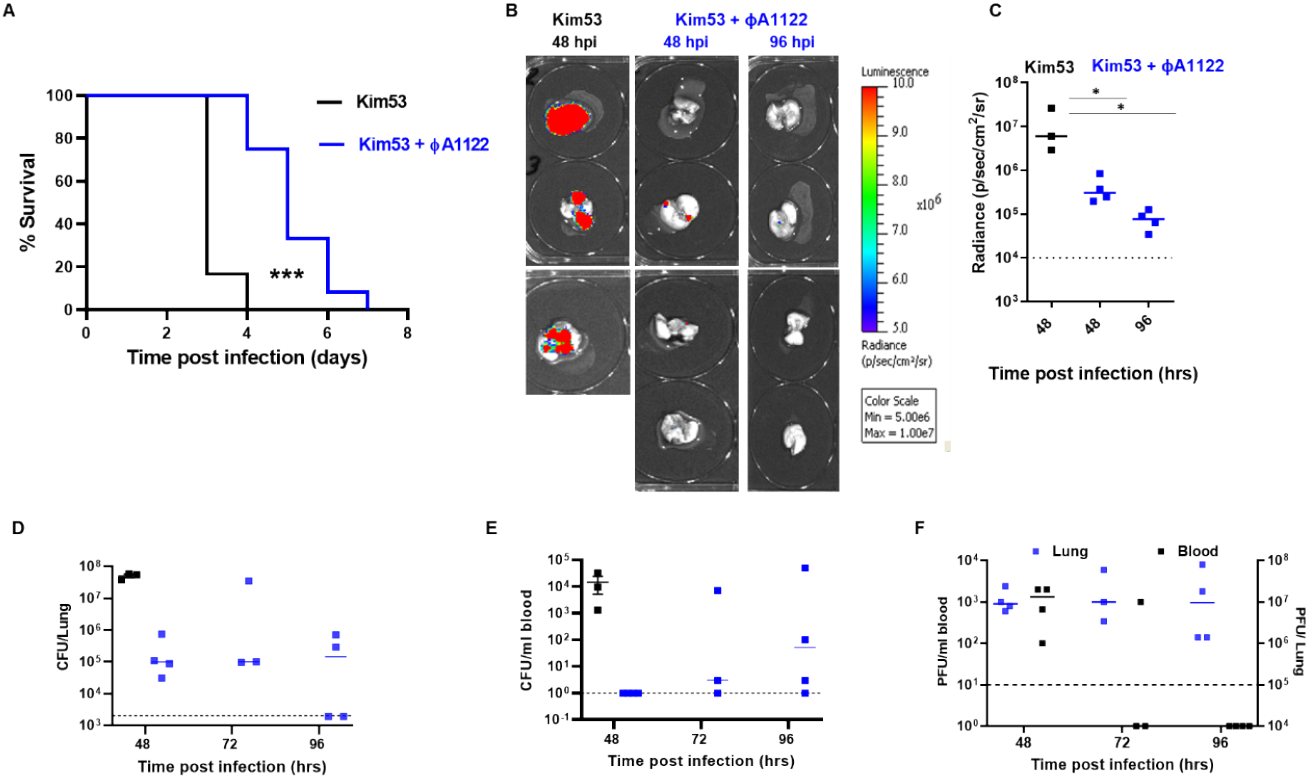
Intranasal administration of a single-dose phage treatment delayed disease progression. C57BL/6J mice were intranasally infected with 10xLD_50_ *Y. pestis* Kim53 strain (A) or Kim53-*lux* luminescent strain (B-F). For phage treatment, one dose of 1×10^9^ ϕA1122 PFU/mouse was IN administered at 5 hpi. Control mice were subjected to intranasal administration of PBS. **A**. Survival curves of control mice (black line, n=6) and phage-treated mice (blue line, n=12). **B-C**. Control (n=3) and phage-treated mice (n=4) were anesthetized at the indicated time points post *Y. pestis* infection. Lungs were harvested, and imaging was performed using IVIS as detailed in the Materials and Methods section. **D-E**. The bacterial load in the lungs and blood was quantified by plating serial dilutions of tissue homogenate/blood on BHIA plates supplemented with 200 μg/ml ampicillin and counting the colonies. **F**. Phage titration was performed by the spot assay technique as detailed in the Materials and Methods section. Each point represents the phage load in the lungs (blue squares, total PFU/lung) or blood (black squares, PFU/ml) of an individual mouse. Horizontal bars represent median values. Dotted lines mark the limit of detection. Statistically significant differences between groups are denoted by asterisks (*, *P* < 0.05; *** *P* < 0.0001, [log rank [Mantel–Cox] test]). Bars indicate standard errors of the means.

To assess *Y. pestis* loads in the lungs of treated mice, we infected mice with a luminescent *Y. pestis* derivative (Kim53-*luxCDABE)*, harvested the lungs at days 2 and 4 pi and visualized bacterial loads using an *in vivo* imaging system (IVIS). We found that at 48 h pi (hpi), lungs from the nontreated control mice showed significantly higher levels of luminescence that those from phage-treated mice (Fig 2, B-C). Bacterial live counts in the lungs aligned with the IVIS results, demonstrating > 2 orders of magnitude higher bacterial loads in control vs. phage-treated mice at 48 hpi (Fig 2D). Similarly, inhibition of *Y. pestis* growth in the lungs of phage-treated mice correlated with high concentrations of ϕA1122 (Figs 2D and 2F, respectively).

Next, we assessed *Y. pestis* dissemination into the blood. During the first 48 h, the bacterial concentration in the blood of the control mice reached ∼ 10^4^ colony-forming units (CFU)/ml, while no bacteria were detected in the blood of phage-treated mice (Fig 2E). However, in the following days, bacteria also appeared in the circulation of phage-treated mice (Fig 2E). The appearance of *Y. pestis* in the blood negatively correlated with the phage amount, which was low at 48 hpi and decreased further to under the limit of detection at 96 hpi (Fig 2F). We therefore concluded that while *Y. pestis* growth in the lungs of treated mice was restricted, the reduction in phage amount in the blood during disease progression enabled *Y. pestis* dissemination/propagation in the circulation, which finally led to mouse death (Fig 2A).

### Multiple-dose phage administrations did not improve treatment efficacy

To improve phage treatment and increase phage load in the blood and internal organs, we designed additional treatment regimens (Fig 3A). Pharmacokinetic analysis showed that in comparison to IN administration, IP injection of ϕA1122 led to higher phage concentrations in the blood, spleen and liver (Fig 1); thus, we added IP injection doses after the initial IN treatment. Moreover, since phages are rapidly cleared from the blood and liver (Fig 1), we provided serial phage treatment in 24-h intervals for 6 days pi. As depicted in Fig 3B, phage treatment was effective in delaying mouse mortality in comparison to that in the nontreated group, as shown above (MTTD = 6 and 3 days, respectively). However, no differences were found among the various phage-treated groups. Thus, increasing phage loads in internal organs during the course of the disease did not improve infection outcomes, suggesting that ϕA1122 lytic activity in the blood might be insufficient.

**Fig 3.**
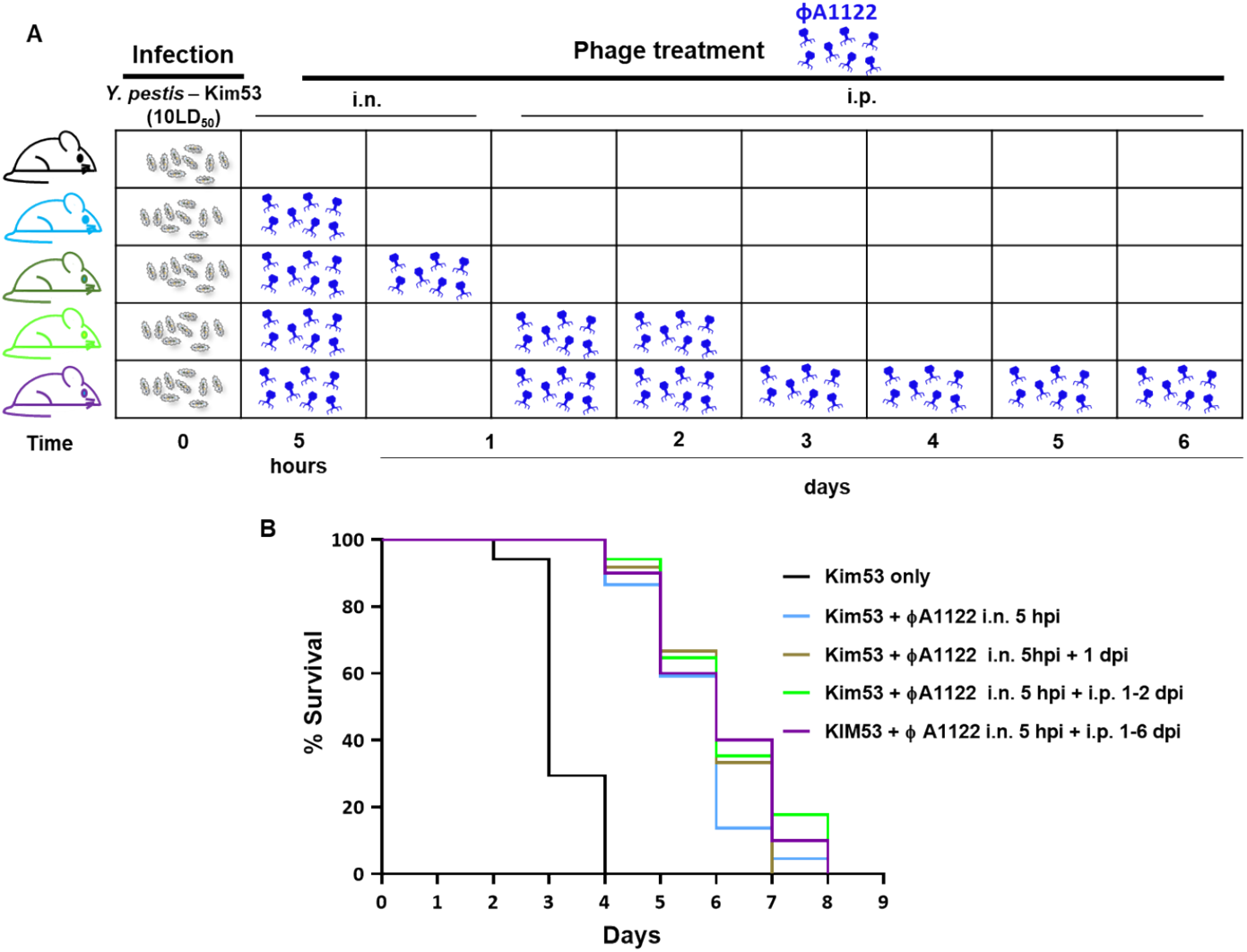
Multiple-dose ϕA1122 phage treatments did not improve treatment efficacy. Schematic representation of mouse treatment regimens (A) and survival curves (B). C57BL/6J mice were IN infected with 10xLD_50_ *Y. pestis* Kim53. For IN phage treatment, mice were inoculated with 35 μl of 1×10^9^ ϕA1122 PFU; IP injection included administration of 0.5 ml of 1×10^9^ ϕA1122 PFU. The treatment regimens were as follows: 1 dose of IN ϕA1122 at 5 hpi (n= 22; light blue line), 2 doses of IN ϕA1122 at 5 hpi and 24 hpi (n=12, olive green line), 1 dose of IN ϕA1122 at 5 hpi + IP injections at 24 + 48 hpi (n=17, green line) and 1 dose of IN ϕA1122 at 5 hpi + IP injections for 6 days, every 24 h (n=10, purple line). Control mice: n=17, black line. B. Statistically significant differences between the control group and phage-treated groups are denoted by asterisks (***, P < 0.0001; log rank [Mantel–Cox] test).

### PST phage shows improved persistence and activity in mouse blood compared to those of ϕA1122

Since the addition of IP administered phage doses did not improve mouse survival (Fig 3B), we tested the possibility that mouse blood may inhibit phage activity. We performed an *in vitro* lysis assay with ϕA1122 and an additional lytic phage, PST. As blood precludes the use of bacterial turbidity as a measure of lysis, we used a luciferase-expressing *Y. pestis* strain as the host and monitored the intensity of the bioluminescent signal in the presence or absence of ϕA1122 or PST. As shown in Fig 4, mouse blood inhibited ϕA1122 lytic activity, whereas the lytic activity of PST was preserved (although to a lesser extent than in that in BHI broth).

**Fig 4.**
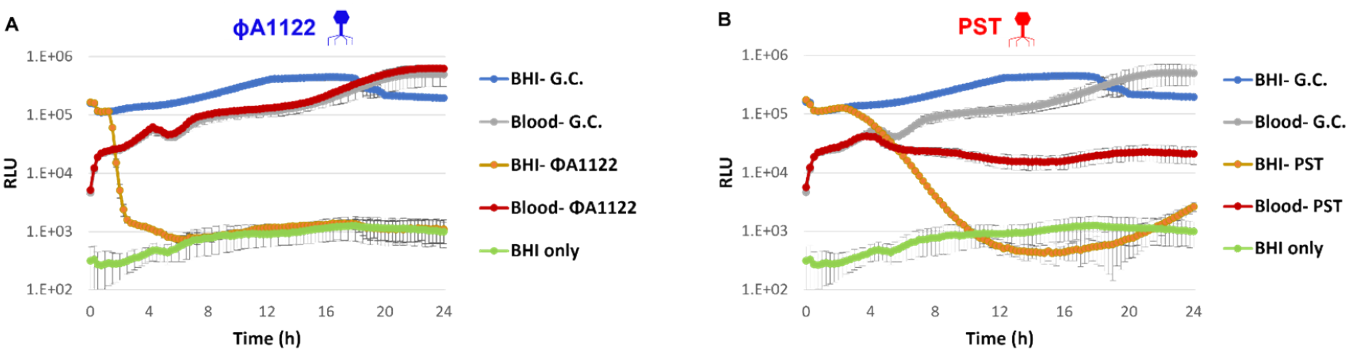
PST is more potent than ϕA1122 in the presence of blood. Phage-based lysis assays were performed with the bioluminescent *Y. pestis* strain EV76-*lux* (10^7^ CFU/ml) suspended in BHI broth (blue and orange lines) or in mouse whole blood (gray and red lines). The *Y. pestis* strain was infected with ϕA1122 or PST phages (10^6^ PFU/ml; multiplicity of infection [MOI] = 0.01). Bioluminescence (RLU) was measured at 37°C in 15 min intervals for 24 h using a Spark 10M plate reader. The experiment was performed in biological duplicates (using blood pooled from 3 mice for each experiment), and the results are representative of one experiment. Values are the average results from 3 wells set up in triplicate in a single experiment, and the error bars represent the standard deviation (SD). The green line represents the background from BHI broth.

As PST was advantageous over ϕA1122 in its lytic activity in blood, we further evaluated its dissemination in mouse tissues following IN or IP administration. As shown in Fig 5, PST disseminated rapidly into mouse tissues following both IN and IP administration, as was observed for ϕA1122 (Fig 1). However, in contrast to the rapid clearance of ϕA1122 from the blood (to under the LOD by 72 h post administration, Fig 1), PST presence in the blood was detected even at 4 days post IN or IP administration.

**Fig 5.**
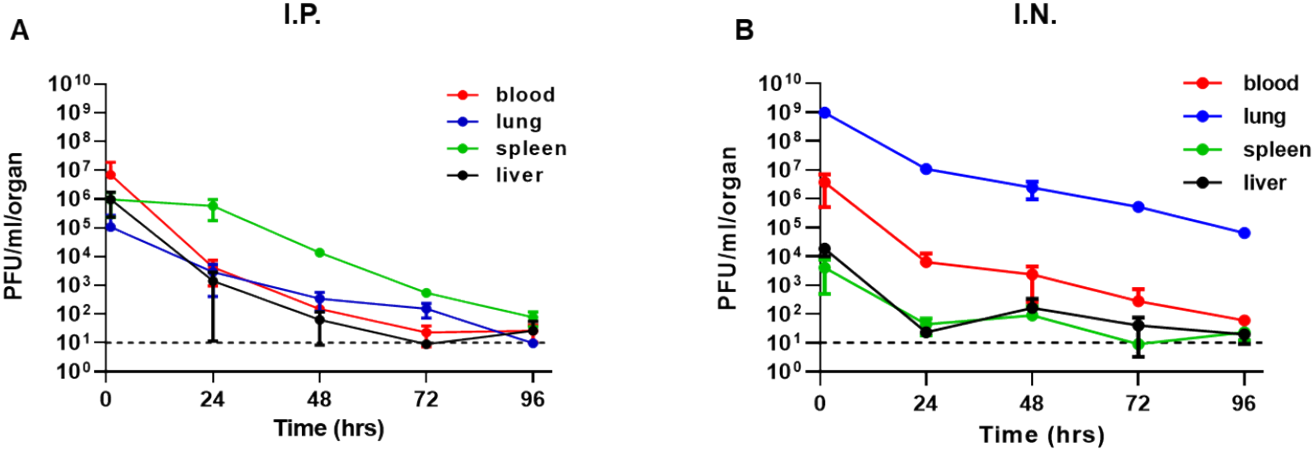
Pharmacokinetic analysis of PST in naïve mice. The experiment was performed as described in the Fig 1 legend.

### Comparing the protective potential of PST and ϕA1122

Next, we compared the therapeutic potential of the phages. The treatment regimen with PST and ϕA1122 is schematically outlined in Fig 6A, and the results indicated that both phages are comparable in their antiplague protective abilities (Fig 6B). This analysis indicates that phage therapy alone is limited in its ability to protect against acute pneumonic plague.

**Fig 6.**
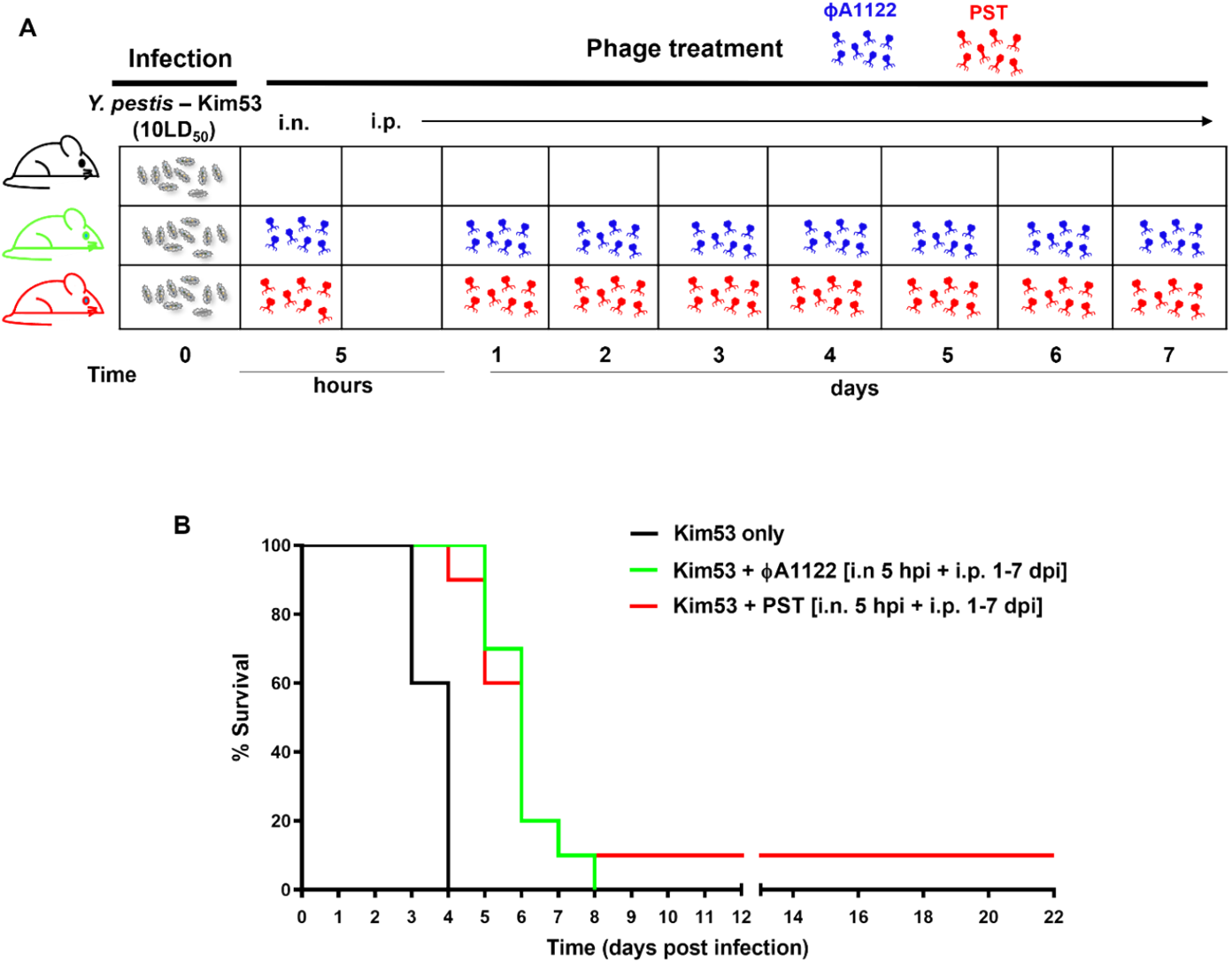
Treatment with ϕA1122 or PST against pneumonic plague has similar outcomes. A. Schematic presentation of the phage treatment regimen. C57BL/6J mice were IN infected with 10xLD_50_ *Y. pestis* Kim53 (gray symbols), followed by PBS (control group) or phage administration. Each dose of PST (red phage symbol) or ϕA1122 (blue phage symbol) suspension contained 1×10^9^ phages. B. Survival curve. The mouse groups were as follows: no phage (n=5, black line), IN phage administration at 5 hpi followed by IP injections every 24 h at days 1-7 post bacterial infection (n=10; green line for ϕA1122 and red line for PST).

### Phage-ceftriaxone combination therapy is highly effective against pneumonic plague

In the event of a plague outbreak involving MDR bacteria, second-line antibiotics may be used as an alternative treatment. Cephalosporins are broad-range antibiotics that are active against *Y. pestis* strains *in vitro* [1], but they are limited in their activity *in vivo* [21,22]. To assess the therapeutic value of adjunctive phage therapy to second-line antibiotic treatment, C57BL/6 mice were IN challenged with a high dose of *Y. pestis* Kim53 (100xLD_50_) and treated with phage, ceftriaxone or a combination of both (Fig 7A). For phage treatment, we used a cocktail composed of both ϕA1122 and PST (1×10^9^ PFU of each per dose). Antibiotic treatment with ceftriaxone rescued only 20% of the mice (Fig 7B), and as was found for treatment with PST or ϕA1122 alone (Fig 6), treatment with the phage cocktail delayed mortality; however, all mice succumbed to the infection (Fig 7B). In contrast, the phage-antibiotic combination treatment was highly effective, leading to the survival of all infected animals (Fig 7B) and clearance of the pathogen from the spleens of all surviving animals at day 21 pi. In addition, surviving animals developed high serum IgG antibody titers against the protective F1 antigen (GMT=25,000 at 21 days pi), implying that the combined treatment also enabled the development of adaptive immunity toward *Y. pestis*.

**Fig 7.**
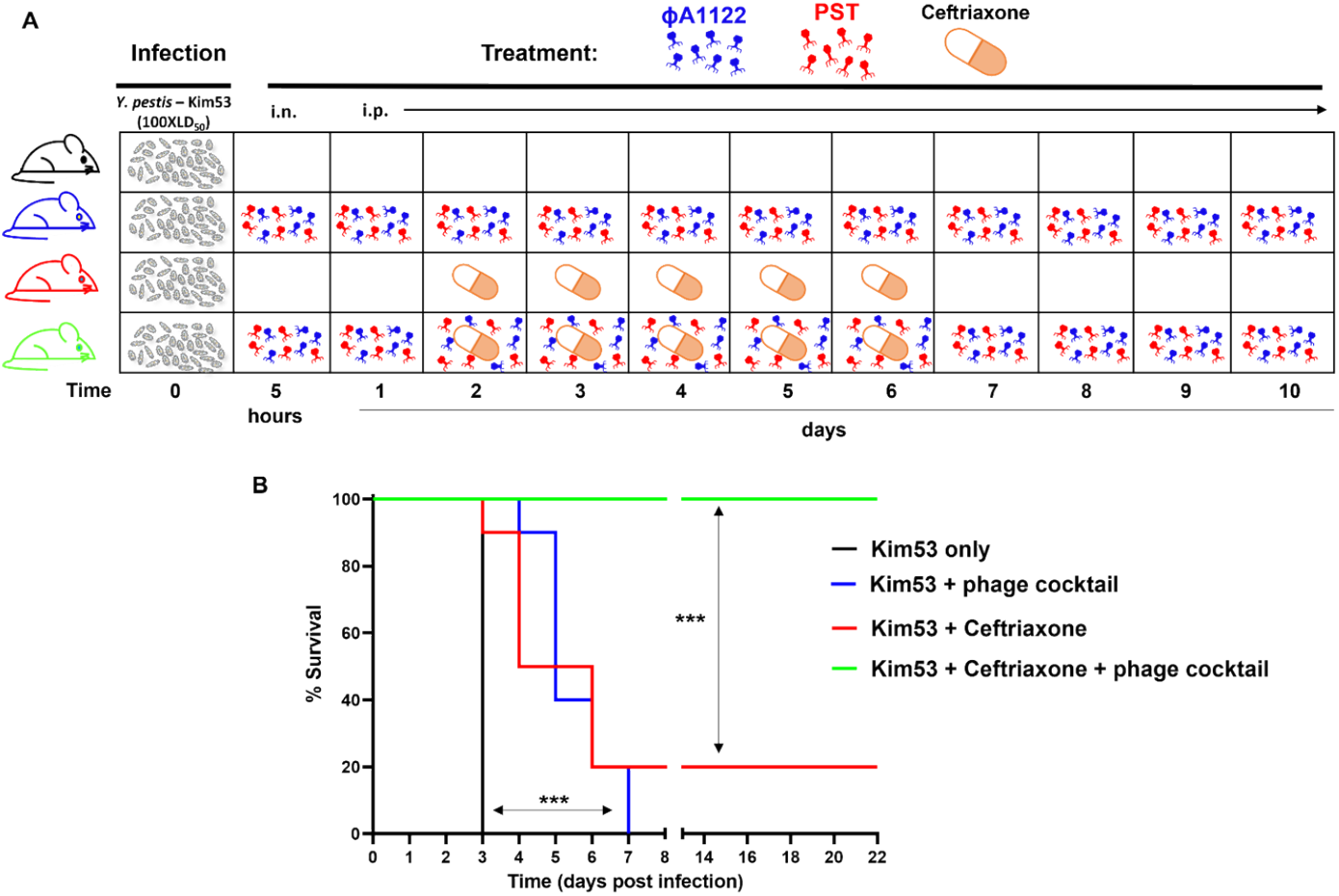
Effective rescue of infected mice by phage-antibiotic combination treatment. C57BL/6J female mice were IN infected with 100xLD_50_ *Y. pestis* Kim53. Mouse groups included control nontreated (n=4), phage-treated (n=10), ceftriaxone-treated (n=10) and phage-ceftriaxone combination-treated mice (n=9). Phage treatment was performed using a phage cocktail composed of ϕA1122 and PST (1×10^9^ each phage/1 dose; 35 μl for intranasal administration or 0.5 ml for IP injection). Treatment included IN administration at 5 hpi followed by IP injections at days 1-10, with 24-h intervals. Ceftriaxone was subcutaneously injected every 12 h on days 2-6 post bacterial infection. Mice were monitored for 22 days. Statistically significant differences are denoted by asterisks (***, P < 0.001; log rank [Mantek-Cox] test).

## Discussion

The experience of the current emerging COVID-19 pandemic with its broad effects on humankind has highlighted the need for preparedness for bacterial-derived outbreaks. In this regard, a major concern is the appearance of highly transmissible MDR pathogens for which current antibiotics may become ineffective. In the case of the plague-causing *Y. pestis* bacterium, antibiotics can effectively treat plague and prevent *Y. pestis* spread in the population. However, a noncontrolled outbreak, which may cause high lethality, may evolve in the case of the emergence and spread of natively derived or deliberately engineered MDR strains. In such circumstances, clinicians will have to use less effective second-line antibiotics for treatment. Adjunctive phage therapy may improve the efficiency of such second-line antibiotics and serve as a safe alternative treatment in the event of MDR infections.

The potential of phages as antiplague therapy was reported almost a century ago (in 1925) by Felix D’Herelle, who injected *Y. pestis*-specific phage preparations into the buboes of four plague patients, all of whom recovered [23]. However, following inconclusive results obtained with additional phage treatment experiments and with the availability of effective antibiotics, interest in developing phage-based therapy against plague decreased. In one of the few studies in which the effectiveness of phage therapy against plague was examined, Filippov and his colleagues assessed the outcome of phage therapy with the ϕA1122 phage in a mouse model of bubonic plague. The study demonstrated that phage treatment was able to delay mouse mortality and provide protection to some of the infected animals [24].

In the present study, we explored the potential of phage therapy (ϕA1122 and PST) alone and in combination with the second-line antibiotic ceftriaxone against pneumonic plague using a mouse model. The *Y. pestis*-specific lytic phages ϕA1122 and PST were chosen for treatment as they are well characterized, and their genomic sequence is known, showing no genes encoding integrase, known antibiotic resistance proteins or toxins [25,26]. The ϕA1122 phage is well suited for use for treatment, as it was shown to be universal against thousands of *Y. pestis* strains and thus is used by the CDC and the U.S. Army Medical Research Institute of Infectious Diseases for *Y. pestis* diagnostics [26,27]. In addition, ϕA1122 and PST bind to different receptors, making their combination as a cocktail a preferable formula to prevent the selection of phage-resistant mutants [25,26].

To evaluate phage therapeutic potential, we infected C57BL/6J mice by intranasal administration of a lethal dose of the *Y. pestis* Kim53 strain followed by treatment with the ϕA1122 and PST phages alone or as a cocktail. IN treatment with only one dose of phage at 5 hpi led to a significant delay in mortality, which correlated with a lower bacterial load in the lungs and slower kinetics of bacterial dissemination/propagation in the blood that those in control nontreated counterparts (Fig 2). However, although disease progression was delayed, phage administration was unable to provide protection, and all treated animals succumbed to the infection (Figs 2-3). Delayed mortality seemed to be associated with low phage concentrations in the circulation (Figs 1 and 5). However, IP administration of additional phage doses to elevate phage titers in the blood did not improve mouse survival in comparison to that with treatment with a single IN dose (Figs 3 and 6). Therefore, the limited treatment efficiency may indicate a reduction in phage lytic activity in the presence of mouse blood (Fig 4), as we have recently observed in human blood [28] and has been reported for other phages [29,30]. However, the PST phage, which retained *in vitro* activity in the presence of blood, still yielded a similarly low therapeutic efficiency *in vivo* as ϕA1122 (Fig 6), indicating that additional limiting factors exist.

It is possible that the presence of ϕA1122 and PST in the spleen and the liver, as reflected by the pharmacokinetic analysis, may indicate phage clearance by the reticuloendothelial system, whereby phages are not free to infect bacteria [31]. Similar observations for ϕA1122-treated bubonic plague mice led Filippov and his colleagues to suggest that the phage and pathogen reside in different compartments in the spleen and liver; therefore, the phage cannot infect and clear the bacterium [24,32]. Finally, the development of phage resistance seems improbable in our model since it has been shown that mutations in the *Y. pestis* lipopolysaccharide (LPS) phage receptors attenuate *Y. pestis* virulence [25] and both ϕA1122 and PST use the LPS receptor to infect the pathogen [25,26].

Although phage treatment failed to save the infected animals, the significantly delayed mortality encouraged us to evaluate the potential of phage as adjunctive therapy to second-line antibiotics as an alternative countermeasure against MDR *Y. pestis*. It has been shown in various case reports and animal model studies that combining antibiotics with phages may enhance bacterial clearance, increase antibiotic penetration into biofilms and decrease the emergence of phage resistance (see reviews [33,34]).

For the combined phage-antibiotic treatment, we used a phage cocktail composed of ϕA1122 and PST, gaining the high observed efficiency of ϕA1122 of *Y. pestis* elimination in the lungs and PST lytic activity in the presence of blood. Additionally, ϕA1122 and PST bind to different sites in the LPS molecule [31,35] and thus targeting different receptors. It is recommended to use phage cocktails composed of several lytic phages targeting various receptors for efficient phage therapy, as this approach broadens the host range and decreases the probability of phage resistance evolution.

For the combined treatment, we used ceftriaxone, an antibiotic that has been shown to have potent *in vitro* activity against *Y. pestis* strains [36] but limited ability to provide protection in a mouse model of pneumonic plague [21,22]. Indeed, when mice were treated with ceftriaxone as a single treatment, 80% of the animals succumbed to the infection (Fig 7B). In contrast, the combined treatment of ceftriaxone and the phage cocktail was highly effective, leading to the survival of all infected animals (Fig 7B) and clearance of the pathogen from internal organs. Surviving animals developed high antibody titers against the protective F1 capsular antigen, indicating the development of adaptive immunity that probably contributed to the combined treatment efficacy.

In conclusion, our results suggest a synergistic effect between phage and antibiotic treatment and highlight the potential of these combinations for emergency use even in cases of acute infections such as pneumonic plague. Notably, training phages to efficiently lyse bacteria in the presence of blood components may improve the therapeutic outcome.

Further research on phage combinations with antibiotics as well as the potential for resensitization to antibiotics may broaden the potential of phage therapy as an alternative therapy against MDR pathogens.

## Materials and methods

### Bacteria, phages and growth media

**Table 1.** *Y. pestis* strains used in the present work.

| Strain | Use | Virulence | Reference |
| --- | --- | --- | --- |
| Kimberley53<br>(Kim53) | Challenge in animal experiments | + | [37] |
| Kimberley53-*luxCDABE*<br>(Kim53-*lux*) | IVIS imaging of bacteria in the lungs | + | [20] |
| Kim53ΔpPCD1ΔpPCP1<br>(Kim53Δ70Δ10) | Phage preparation and titration | - | [37] |
| EV76-*luxCDABE*<br>(EV76-*lux*) | Phage lysis assay | - | [28] |

The Kim53 and Kim53Δ70Δ10 strains were grown on Brain-Heart-Infusion Agar (BHIA; 241830, BD, USA), and Kim53-*lux* and EV76-*lux* were routinely grown on BHIA supplemented with ampicillin (200 μg/ml, A0166, Sigma–Aldrich, Israel).

The *Y. pestis*-specific lytic phages used in this study were ϕA1122 (accession no. NC004777, kindly provided by professor Mikael Skurnik [38]), and PST (ATCC, cat. no.: 23207-B1; accession no. KF208315).

### Animal studies

The study was performed in accordance with the Recommendations for the Care and Use of Laboratory Animals (National Institutes of Health [NIH]). Animal experiments were performed in accordance with Israeli law and were approved by the Institutional Ethics Committee for animal experiments (protocols no.: M-68-17, M-59-18, M-47-19, M-01-20, M-72-20 and M-25-21). Experiments were performed in an animal biosafety level 3 (ABSL-3) laboratory. Female 8-to 12-week-old mice (C57BL/6J; Invigo, Israel) were used for all animal experiments.

For IN challenge, bacterial colonies of Kim53 or Kim53-*luxCDABE* were grown on heart infusion broth (HIB, 238400, BD, USA) supplemented with 0.2% xylose and 2.5 mM CaCl_2_ (Sigma–Aldrich, Israel) at 28°C for 22 h, as previously described [20]. Counting of CFU was performed by plating 0.1 ml of the appropriate culture dilutions on BHIA plates and incubating them for 48 h at 28°C. Prior to infection, mice were anesthetized with a mixture of 0.5% ketamine HCl and 0.1% xylazine and then infected IN with 35 μl of the bacterial suspension containing 10xLD_50_ or 100xLD_50_ Kim53 (1xINLD_50_ = 1,100 CFU [39]) per mouse.

For monitoring of bacterial dissemination to the lungs and blood, mice were anesthetized. Blood was collected by cardiac puncture, and lungs were harvested and plated on BHIA supplemented with streptomycin (100 μg/ml, S6501, Sigma–Aldrich, Israel).

For phage pharmacokinetic analysis, one dose of ϕA1122 or PST phage solution (1×10^9^ PFU/mouse) was IN administered to C57BL/6J mice (35 μl) or delivered by IP injection (0.5 ml). Mice were anesthetized prior to IN administration. For phage enumeration in organs, mice (n=3 per time point and per administration route) were sacrificed, and blood was collected by cardiac puncture followed by serial 10-fold dilution in SM buffer (0.1 M NaCl, 8 mM MgSO_4_, 50 mM Tris-HCl pH 7.5 and 0.01% gelatin solution). The lungs, spleen and liver were harvested, washed in PBS and transferred to 1 well in 6-well microplates. The spleen, liver and lungs were crushed in PBS (1 ml for spleen and liver and 2 ml for lungs). The lung extract was filtered through a 70 μm cell strainer. The crushed tissues were serially diluted 10-fold in SM buffer. Phage titration of the diluted samples was performed by the spot assay technique by dropping 3 drops of 10 μl samples on Kim53Δ70Δ10 lawns grown on BHIA top agar containing 200 μg/ml streptomycin to prevent bacterial contamination [30].

For bioluminescence imaging analysis of luciferase-expressing *Y. pestis*, mice were IN exposed to 10xLD_50_ Kim53-*lux* followed by IN administration of ϕA1122 or PBS (control group). Photon emission from the lungs was visualized using an IVIS (Caliper Life Sciences, Hopkinton, MA). Image acquisition was performed using the following settings: binning of 2 and acquisition times of 1–4 min. The luminescence signals for all images were normalized and reported as photons/second/cm^2^/sr using Living Image® 4 software.

For titration of anti-F1 IgG in serum, blood was collected from mice via the tail vein at day 21 pi, and titers of IgG against F1 were determined by ELISA as previously described [40].

### Phage treatment

Phages were applied as a single suspension (35 μl containing 10^9^ PFU of ϕA1122 or PST) or as a cocktail (ϕA1122 + PST, 10^9^ PFU each). IN administration was conducted 5 h post *Y. pestis* infection. Phage IP injections (0.5 ml, 1×10^9^ PFU/mouse) were conducted at 24 h intervals (regimens described in the relevant figure legend). Prior to phage IN administration, mice were anesthetized by subcutaneous injection of 0.5% ketamine HCl and 0.1% xylazine mixed solution. Control mice were administered PBS.

### Antibiotic treatment

Ceftriaxone (PHR-1382, Sigma–Aldrich, Israel) was injected subcutaneously twice per day for 5 consecutive days, starting at 48 h post *Y. pestis* infection.

### Statistical analysis

The results were analyzed by GraphPad Prism 8.2.0. Pairwise comparisons of mouse groups were determined using a log rank (Mantel–Cox) test. A P value of 0.05 was characterized as the significance threshold.

### Bioluminescence-based lysis assay

Blood from 3 naive C57BL/6J mice was pooled in citrate-containing tubes (Vacutainer sodium citrate tubes, BD, USA). The *Y. pestis* EV76-*lux strain* was grown on BHIA at 37°C for 48 h. Bacterial colonies were suspended in PBS and inoculated (1:10; vol:vol) in BHI broth or in mouse whole blood and transferred (90 μl/well) into a 96-well transparent-bottom white microplate (Thermo Scientific Nunc: cat. no. 165306). Infection was performed by adding 10 μl of ϕA1122 or PST phage solution (MOI = 0.01) or 10 μl of SM buffer to the growth control wells. The bacterial growth curves were assessed by tracking the bioluminescent signal (relative light units, RLU) of each well at 15 min intervals over 24 h using a SPARK 10M plate reader (Tecan). The temperature in all experiments was 37°C.

### Phage preparation and purification

Purified bacteriophage stocks (ϕA1122 or PST phages) were prepared from phage-infected cultures of nonvirulent *Y. pestis* Kim53Δ70Δ10. Phage lysates were prepared by growing bacterial cultures in two 2 l Erlenmeyer flasks containing 500 ml of BHI inoculated with overnight starter culture to obtain an initial OD_600_ of 0.05. Bacterial cultures were incubated at 200 rpm at 28°C until OD_660_=1. Phage solution was added to the bacterial culture (MOI=0.01), and the culture was further incubated under the same conditions for another 4 h for ϕA1122 or 24 h for PST. Culture lysates were centrifuged at 6,000 rpm for 10 min at 4°C followed by 0.45 μm filtration for the removal of residual bacteria or bacterial debris. Bacterial DNA was removed by endonuclease digestion using DENARASE endonuclease (c-LEcta, Germany). Digestion was performed with 20 U/ml endonuclease in the presence of 2 mM MgCl_2_ (Spectrum, New Brunswick, NJ) for 24 h at 4°C under constant mild agitation. Sucrose (St. Louis, MO) was added to the phage solution to a final concentration of 4%. Ultrafiltration was performed using a 300 or 750 kDa (for ϕA1122 or PST purification, respectively) nominal molecular weight cutoff (NMWC) with a 1 mm diameter polyethersulfone (PES) hollow fiber membrane cartridge (Cytiva, Marlborough, MA). Using an ultrafiltration system, the phage solution was concentrated 10-fold by volume and diafiltrated X5 against PBS (Biological Industries, Israel) supplemented with 4% sucrose. The ultrafiltrated phage solution was filtered again through a 0.45 μm filter. Endotoxin removal was performed twice by using a Toxin Eraser endotoxin removal kit (GenScript, Piscataway, NJ) according to the manufacturer’s guidelines. Elution was performed with a PBS-0.4% sucrose solution buffer followed by 0.2 μm filtration. Residual endotoxins in the purified phage preparations were determined by the Limulus Amebocyte Lysate (LAL) Kinetic-QCL Kit (Lonza, Basel, Switzerland) according to the manufacturer’s instructions. Purified phage preparations containing endotoxin units (EU)/1E09 PFU ≤ 2 were used for the phage therapy experiments. The phage stock was stored at 4°C in the dark until use.

## Acknowledgments

We would like to thank Professor Mikael Skurnik for kindly providing the ϕA1122 phage.

